# Unsupervised deconvolution of molecular heterogeneity uncovers novel signatures and glia-neuron ratio

**DOI:** 10.1101/435677

**Authors:** Lulu Chen, Niya Wang, Chunyu Liu, David M. Herrington, Robert Clarke, Zhen Zhang, Yue Wang

## Introduction

While the two major types of cells in the brain are known to be glia and neuron, the true ratio of glia to neurons in the brain remains a mystery. One of recent studies using efficient cell counting method provides compelling evidence for 1:1 ratio on four whole human brains ^1^. The same study also reveals that the ratio of glia to neurons in the brain varies from one region to another, sometimes dramatically, e.g., 3.76:1 in the cerebral cortex versus 1:4.3 in the cerebellum ^1,2^. However, other scientists have argued that more rigorous studies are needed in which just about every known or unknown marker for both neurons and glia is used to capture as many of the different cell types as possible.

Here we report a study that applies UNsupervised DecOnvolution version 2 (UNDO2.0) software tool ^3^, the spin-off from the Convex Analysis of Mixtures (CAM) framework for deconvolving mixtures of more than two subtypes ^4^, to the gene expression and DNA methylation data acquired from heterogeneous brain tissues. UNDO2.0 can automatically detect cell-specific markers located on the scatter radii of mixed gene expressions, estimate cellular proportions in each sample, and deconvolve mixed expressions into cell-specific expression profiles. The in-silico analysis uncovers many novel marker genes and provides the molecularly-defined 2.2:1 ratio of glia to neurons in the brain.

An R package of UNDO2.0 is available at http://bioconductor.org/packages/UNDO.

An R package of CAM is available at http://bioconductor.org/packages/CAMTHC.

## Results

### Re-validation of UNDO2.0 on benchmark datasets

We first test UNDO2.0 on the gene expression data acquired from Rat neuron-astrocyte mixtures (GSE19380 ^5^). We apply UNDO2.0 to the 3 mixtures of neuron and astrocyte, samples 17, 19, 21. Experimental result shows accurate estimate of both cellular proportions (correlation coefficient of 0.997~0.998 between known proportions and the estimates by UNDO2.0) and cell-specific expression profiles (correlation coefficient of 0.984~0.988 between the UNDO2.0 estimates and pure neuron/glia expressions).

We then test UNDO2.0 on the DNA methylation data acquired from post mortem frontal cortex (GSE41826 ^6,7^). The platform is Illumina HumanMethylation450 BeadChip. There are 480,492 probes in total. We apply UNDO2.0 to the 9 reconstituted mixtures of FACS-purified neuron and glia with neuron proportions varying from 10% to 90%. The experimental results are summarized in **Figure 1**. It can be clearly seen that the scatter plot of two mixtures form a parallelogram, where the orange lines represent the two column vectors of the predefined proportions, tightly embraced the scatter distribution. Experimental result shows accurate estimate of cellular proportions (correlation coefficient of 0.998 between the UNDO2.0 estimates and known proportions) given in **Table 1**.

**Figure 1.**
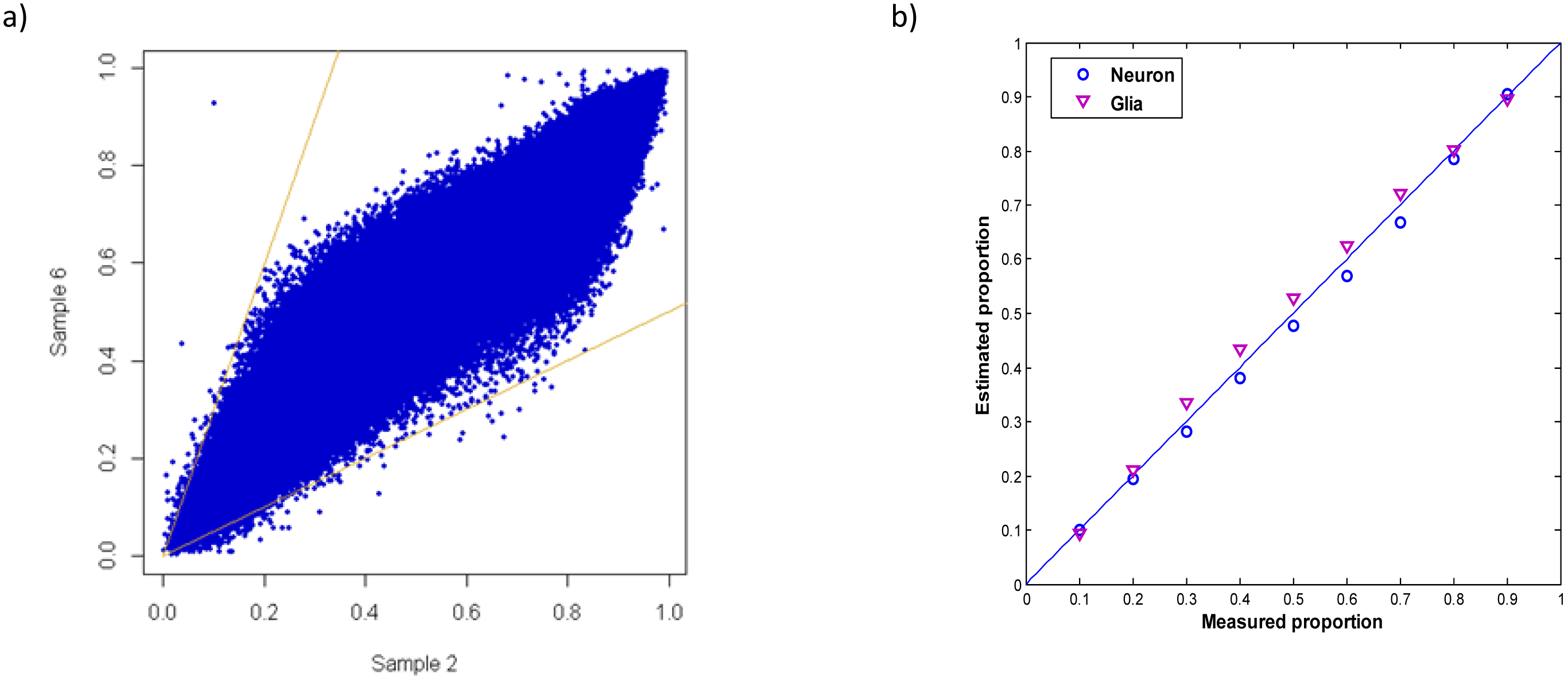
a) Scatter plot of methylation signals in two mixtures. b) Scatter plot of the measured and UNDO2.0 estimated neuron-glia proportions.

**Table 1.**

| Sample # | 1 | 2 | 3 | 4 | 5 | 6 | 7 | 8 | 9 |
| --- | --- | --- | --- | --- | --- | --- | --- | --- | --- |
| Neuron (ground truth %) | 0.1 | 0.2 | 0.3 | 0.4 | 0.5 | 0.6 | 0.7 | 0.8 | 0.9 |
| Neuron (UNDO2.0 estimated %) | 0.101 | 0.195 | 0.281 | 0.381 | 0.476 | 0.568 | 0.667 | 0.785 | 0.906 |

### UNDO2.0 Re-analysis on gene expression data of Huntington’s disease

The gene expression dataset contains 56 samples of caudate nucleus, among which 29 samples are controls and 27 are cases (GSE3790 ^5^). The cases consist of 12 and 15 subjects with grade 1 and 2 Huntington’s disease, respectively. One of the pathological hallmarks on Huntington’s disease is neuronal death and glial proliferation of caudate nucleus ^5^. We apply UNDO2.0 to these three sample groups separately, and estimate both the neuron/glia proportions and neuron/glia-specific expression profiles. The experimental results are consistent with what previously reported by supervised deconvolution method on the same dataset ^5^, as summarized in **Table 2**.

**Table 2.** The means of neuron/glia-specific gene expressions in the three sample groups estimated by UNDO2.0. The average of neuron/glia proportions in the three sample groups estimated by UNDO2.0.

|  | Mean expressions |  | Average proportions |  |
| --- | --- | --- | --- | --- |
|  | Neuron | Glia | Neuron | Glia |
| Control | 427.21 | 173.50 | 0.53 | 0.47 |
| Grade 1 | 386.92 | 257.84 | 0.47 | 0.53 |
| Grade 2 | 380.04 | 262.77 | 0.45 | 0.55 |

Specifically, in Huntington’s disease groups, both mean expression levels and proportions in the neurons decrease, while the opposite changes are observed in glial cells. Furthermore, differential analysis on the neuron/glia-specific expression profiles reveals 496 probe sets significantly up-regulated in neuron and 774 probe sets significantly up-regulated in glia between grade 1 and control; and 294 probe sets significantly up-regulated in neuron and 1082 probe sets significantly up-regulated in glia between grade 2 and control. Moreover, the Ingenuity Pathway Analysis (IPA), focused on canonical pathways using these phenotype-upregulated genes, reconfirms that Huntington’s disease signaling pathways are among the top disease-related pathways (**Appendix**). The experimental results are summarized in **Figure 2**. Lastly, the PPM1H expression levels in neurons estimated by UNDO2.0 shows significant increase in Huntington’s disease compared to control, consistent with what validated by immunohistochemistry method ^5^. PPM1H (NERPP-2C) has been shown to have a developmentally regulated expression and to be expressed by adult neurons in rat.

**Figure 2.**
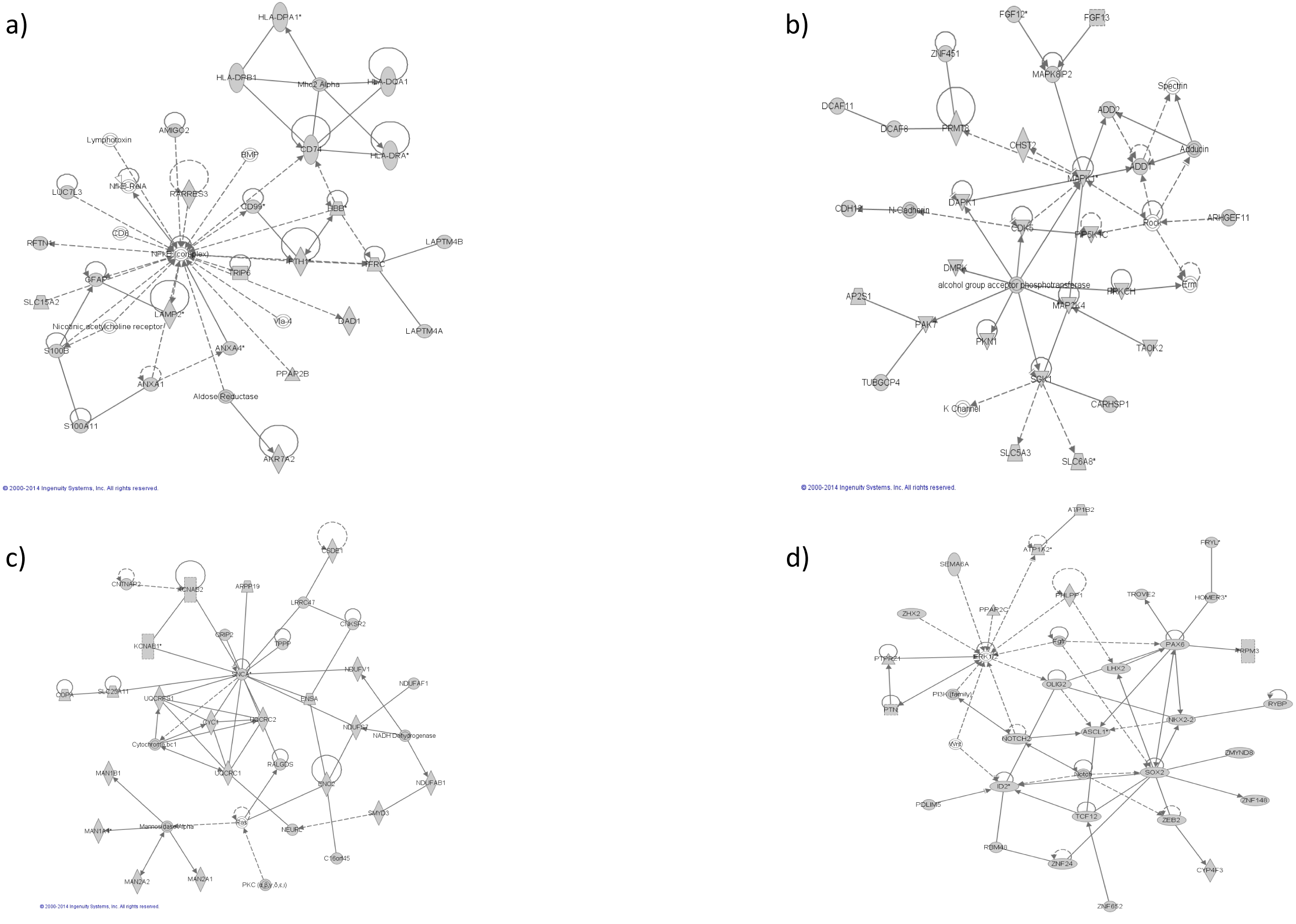
a) One of the top pathway networks enriched by 496 neuron-upregulated probe sets in grade 1. b) One of the top pathway networks enriched by 774 glia-upregulated probe sets in grade 1. c) One of the top pathway networks enriched by 294 neuron-upregulated probe sets in grade 2. d) One of the top pathway networks enriched by 1082 glia-upregulated probe sets in grade 2.

To illustrate the invalidity of the common assumption that the expression signatures of cell-type specific markers are invariant across phenotypes, made necessarily by most supervised deconvolution methods ^5^, we also check the mean expression levels of cellspecific marker genes across the three phenotypic groups, and observe significance changes in the marker gene expressions associated with both neurons (5 genes) and glia (9 genes), shown in **Figure 3**.

**Figure 3.**
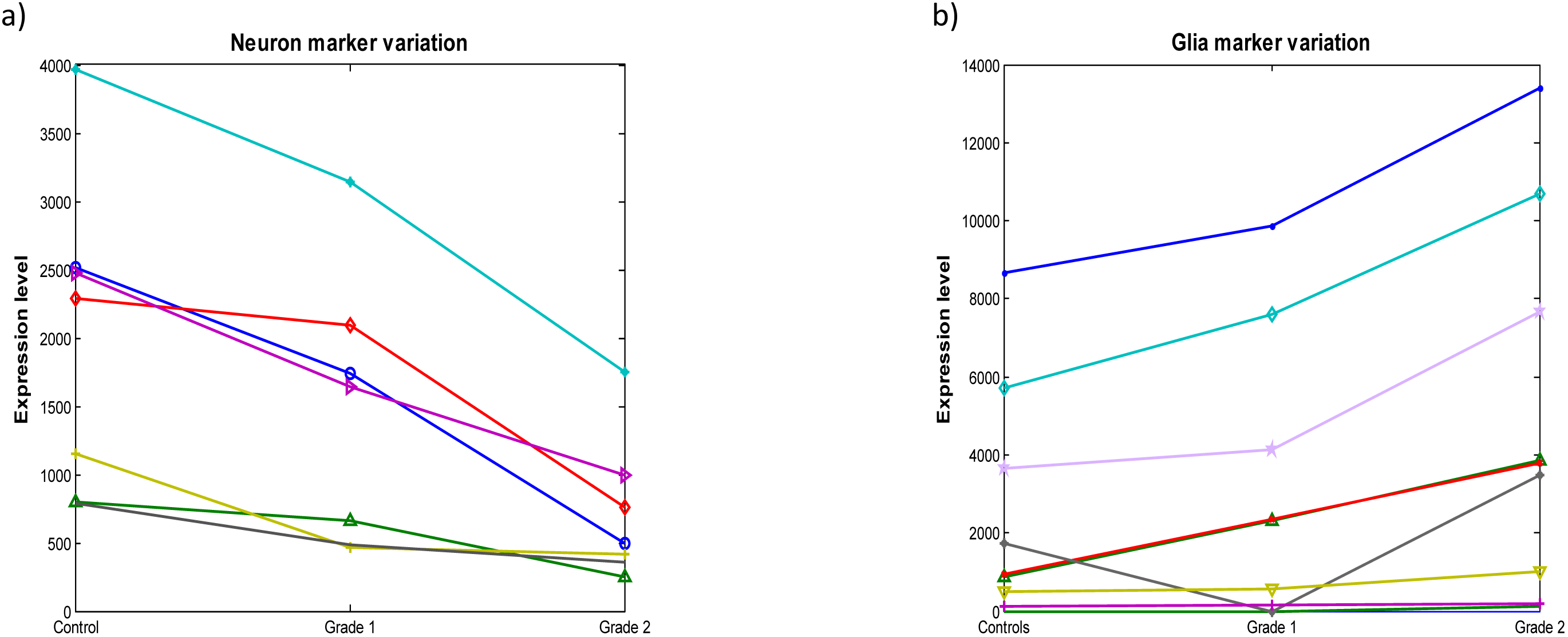
Significant variations in marker gene expression levels (neuron and glia) among control, grade 1, and grade 2.

### UNDO2.0 Analysis on gene expression data of different brain regions

We first apply UNDO2.0 to the gene expression data of 129 samples acquired from cerebellum (CB). The deconvolution results indicate that the overall neuron proportion is about 70% and non-neuron proportion about 30%, consistent with what previously reported ^1,2^. We then apply UNDO2.0 to the gene expression data of 129 samples acquired from parietal cortex (PC). The neuro/glia marker genes blindly detected by UNDO2.0 are geometrically close to (overlap), in the scatter simplex, the marker genes given by literatures ^5^, as shown in **Figure 4**. More interestingly, the deconvolution also reveals that the overall glia to neuron ratios are different between CB and PC, indicating that neuron/glia distributions may be region-dependent.

**Figure 4.**
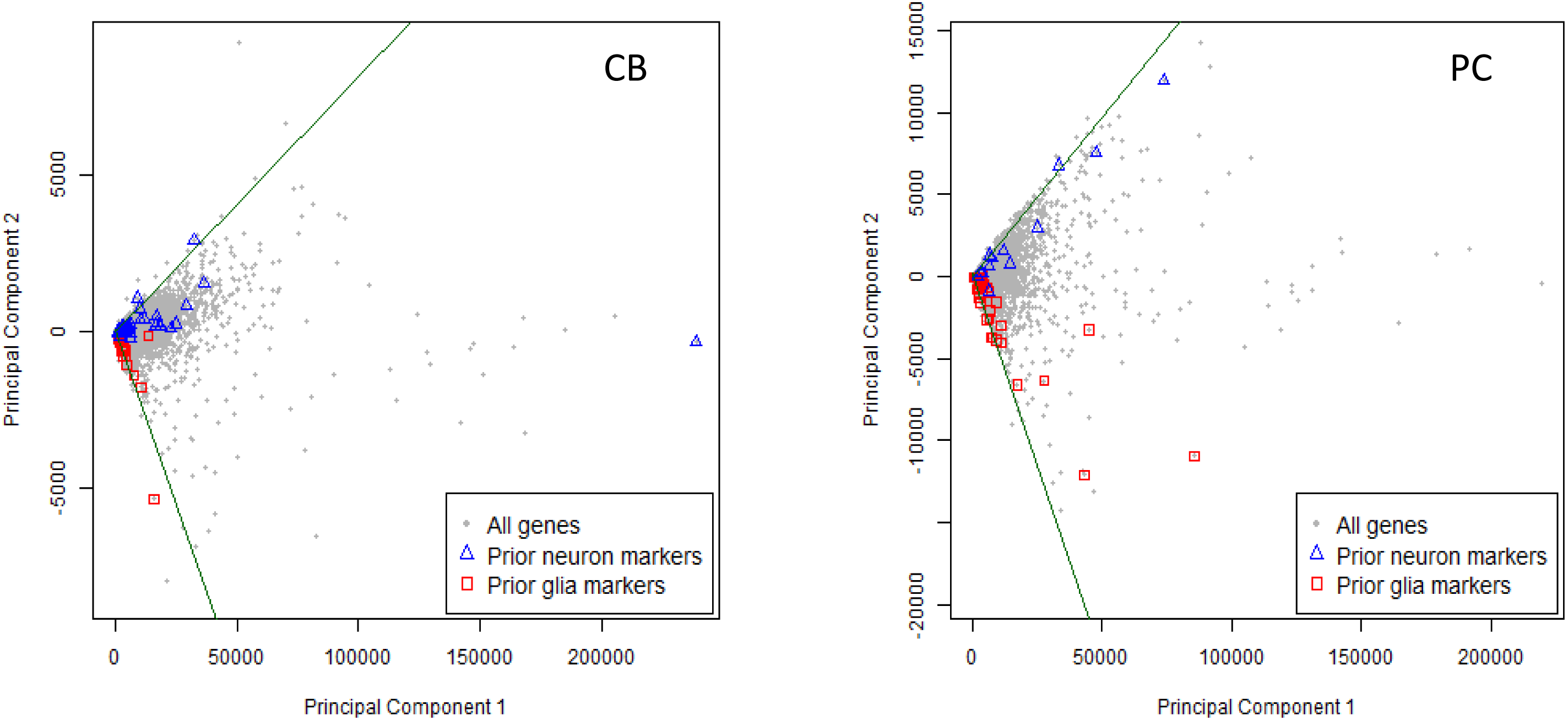
Convex cone in scatter space identified by UNDO2.0, where the marker genes given by literatures are geometrically close to the edges of convex cone.

We then apply UNDO2.0 to the paired 258 samples all together, and detect the two dominant yet most differential neuron cell types that are concordantly associated with CB versus PC brain regions. Strikingly, our experiment detects many region-specific neuron markers that are exclusively enriched in CB or PC neurons, shown in **Figure 5**, molecularly defining two region-dependent neuron subpopulations.

**Figure 5.**
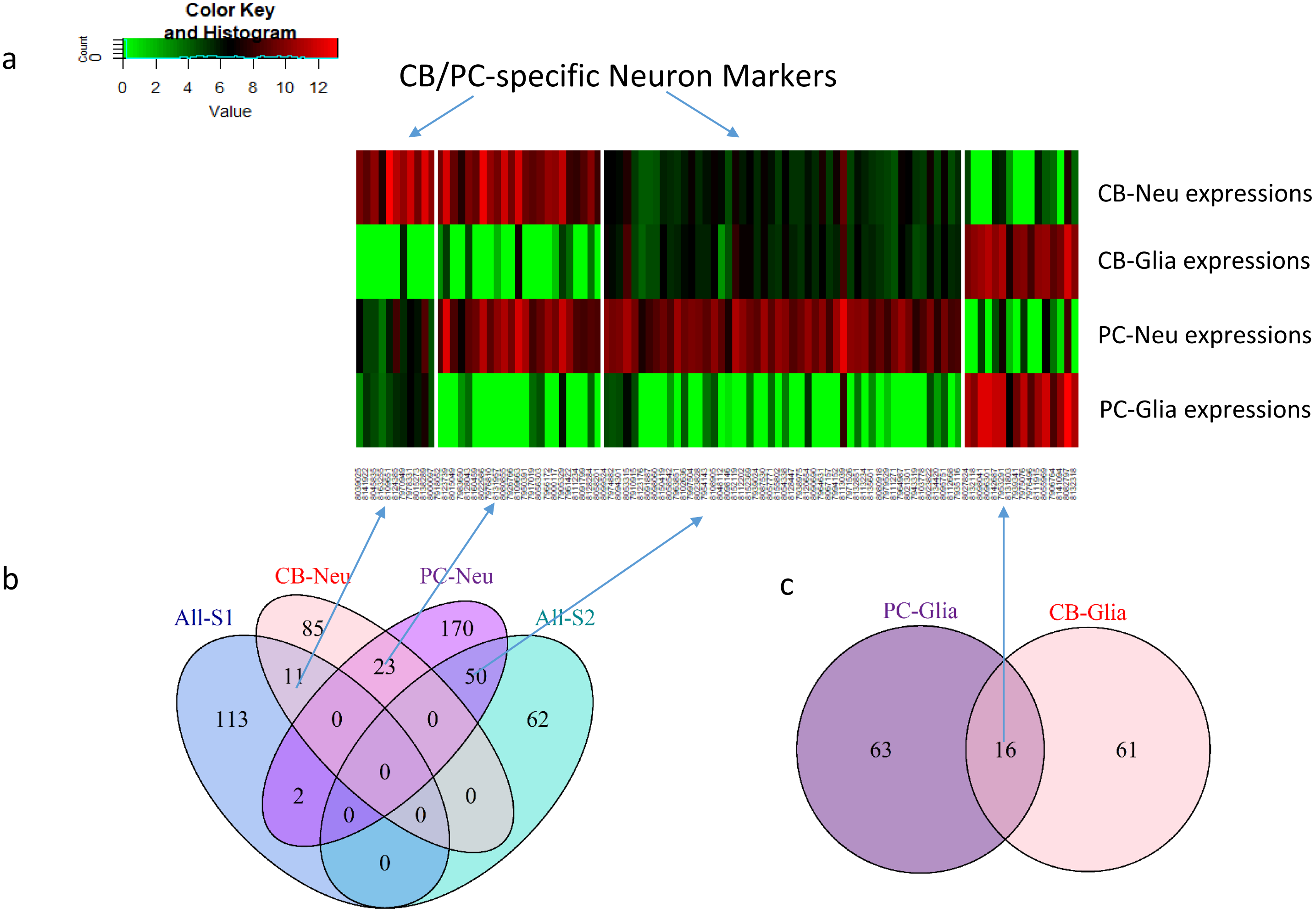
Heatmap of neuron/glia-specific expressions in CB and PC (region-dependent/independent marker genes). (b) Overlap of neuron markers (FC > 20) blindly detected in CB samples, PC samples, and all samples. (c) Overlap of glia markers (FC > 20) blindly detected in CB samples, and PC samples.

### UNDO2.0 Analysis on methylation data in human brain cerebellum region

We apply UNDO2.0 to the methylation data of 130 CB samples and detect two cell types with relative proportions of 30% and 70%, respectively. After a sample-wise comparison on the relative proportion vectors (non-neuron: neuron), estimated using methylation data and that of gene expression data acquired from the same regional samples, we observed a high correlation coefficient of 0.948 (see **Figure 6**) that indicates that the two cellular subpopulations deconvolved from methylation data correspond to neuron and non-neuron cells.

**Figure 6.**
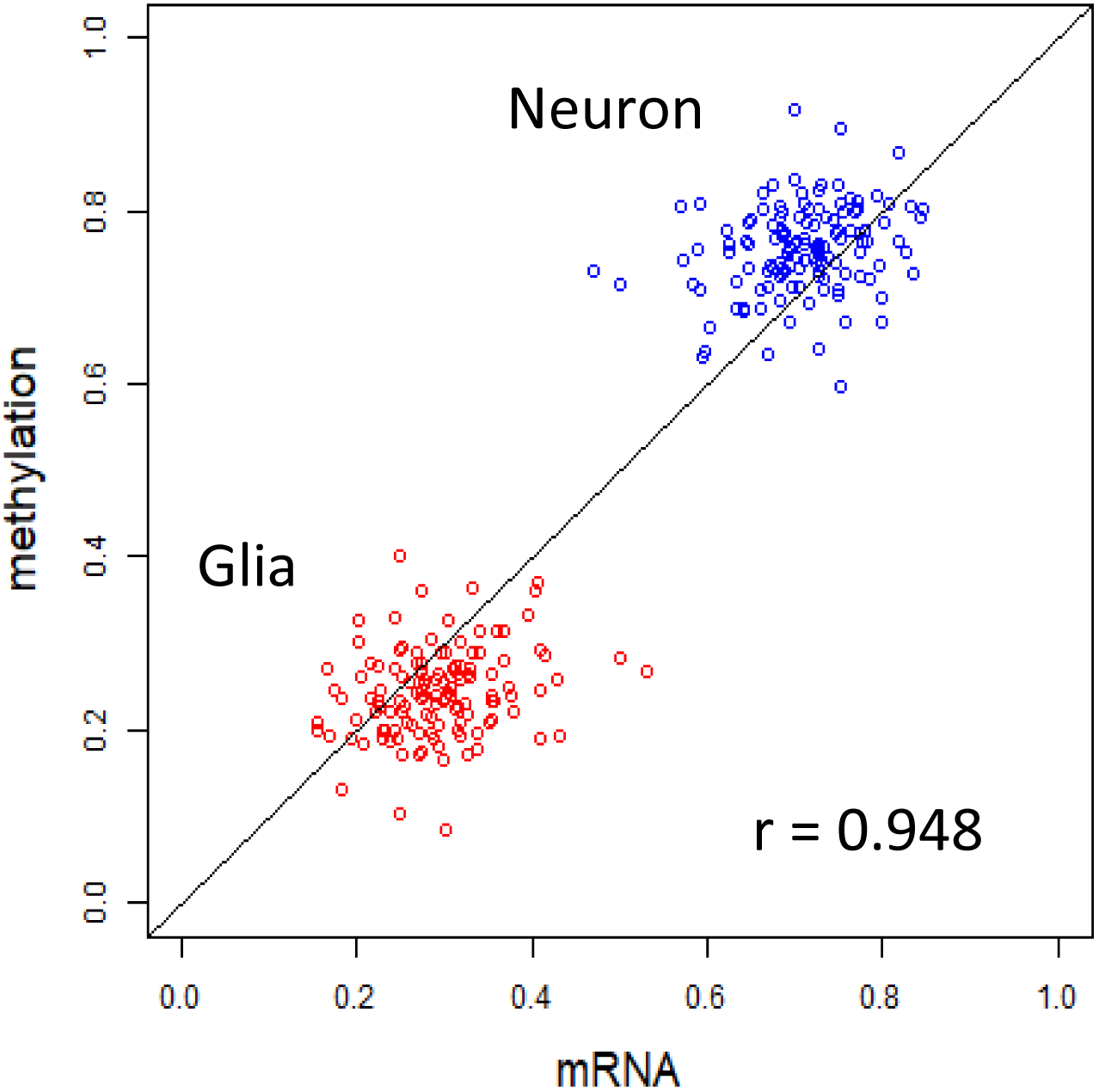
Scatter plot of the relative neuron/non-neuron proportion vectors estimated by UNDO2.0 on methylation data versus gene expression data.

## Discussion

UNDO2.0 uses various molecular signals, e.g., gene expression profiles, and thus detects molecularly define subpopulations. To characterize biological associations of the subpopulations (e.g., cell or tissue types), subsequent enrichment analysis is often required. Because the deconvolution is limited to only two subpopulations, UNDO2.0 produces the two most differential subpopulations, while not necessarily the most dominant ones. Our experimental results show that preprocessing may have significant impact on deconvolution. For example, when sample clustering is performed prior UNDO2.0, the input mixtures would represent dominant subpopulations; while extracted mixtures by e.g. principle component analysis may favor the most differential subpopulations. Furthermore, feature selection (e.g., genes or proteins) based on a priori would produce the subpopulations that may be focused on certain biological questions.

Another important yet subtle concept is the difference between differentially expressed genes (DEGs) and marker genes (MGs). MGs are the genes exclusively enriched in particular subpopulations in the form of e.g. (High, 0, 0); (0, High, 0); or (0, 0, High); where ‘0’ (or near zero) is a much stronger requirement than ‘Low’.

While correlation coefficient is a classic concept, its calculation in comparing the estimates by two different methods needs special care. If the cellular proportions associated with each sample, in a population, are largely random over the entire space of ‘proportion vectors’, then, the correction coefficient between two estimates can be calculated over either one individual component or all components of the proportion vector. In contrast, if such cellular proportions are cluster-structured with moderate randomness’, e.g., the cellular proportions of neuron and glia cells follow an average ratio, then correlation coefficient between two estimates should be calculated using all components of the proportion vector, and within-cluster high correlation may not be expected.

## Methods

See reference ^3^.

### Availability of UNDO2.0 software

The R package of UNDO2.0 is available for academic and non-commercial use at: http://www.bioconductor.org/packages/release/bioc/html/UNDO2.0.html

## ADDITIONAL INFORMATION

Supplementary information accompanies this paper at http://www.

## ACKNOWLEDGMENTS

This work was funded in part by the National Institutes of Health under Grants HL111362-05A1, HL133932, BC171885P1, and U24CA160036-05S1.

## AUTHOR CONTRIBUTIONS

LC and NW developed the UNDO2.02.0 algorithm and performed data analyses; CL, DMH, RC and ZZ provided data and interpreted results; YW designed the framework and wrote the manuscript.

## Appendix

### Grade 1 group (neuron 496 probe sets)

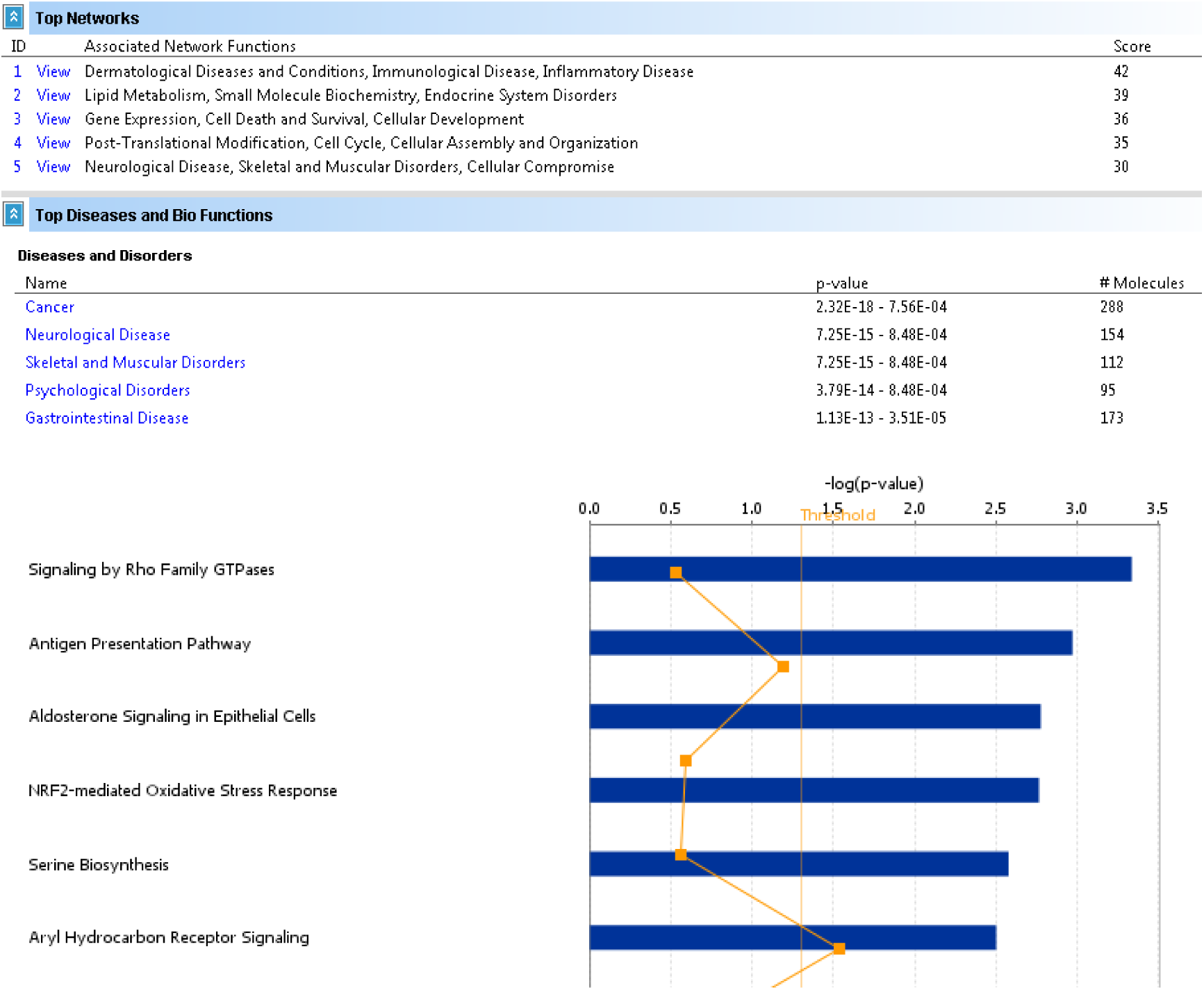

### Grade 1 group (glia 774 probe sets)

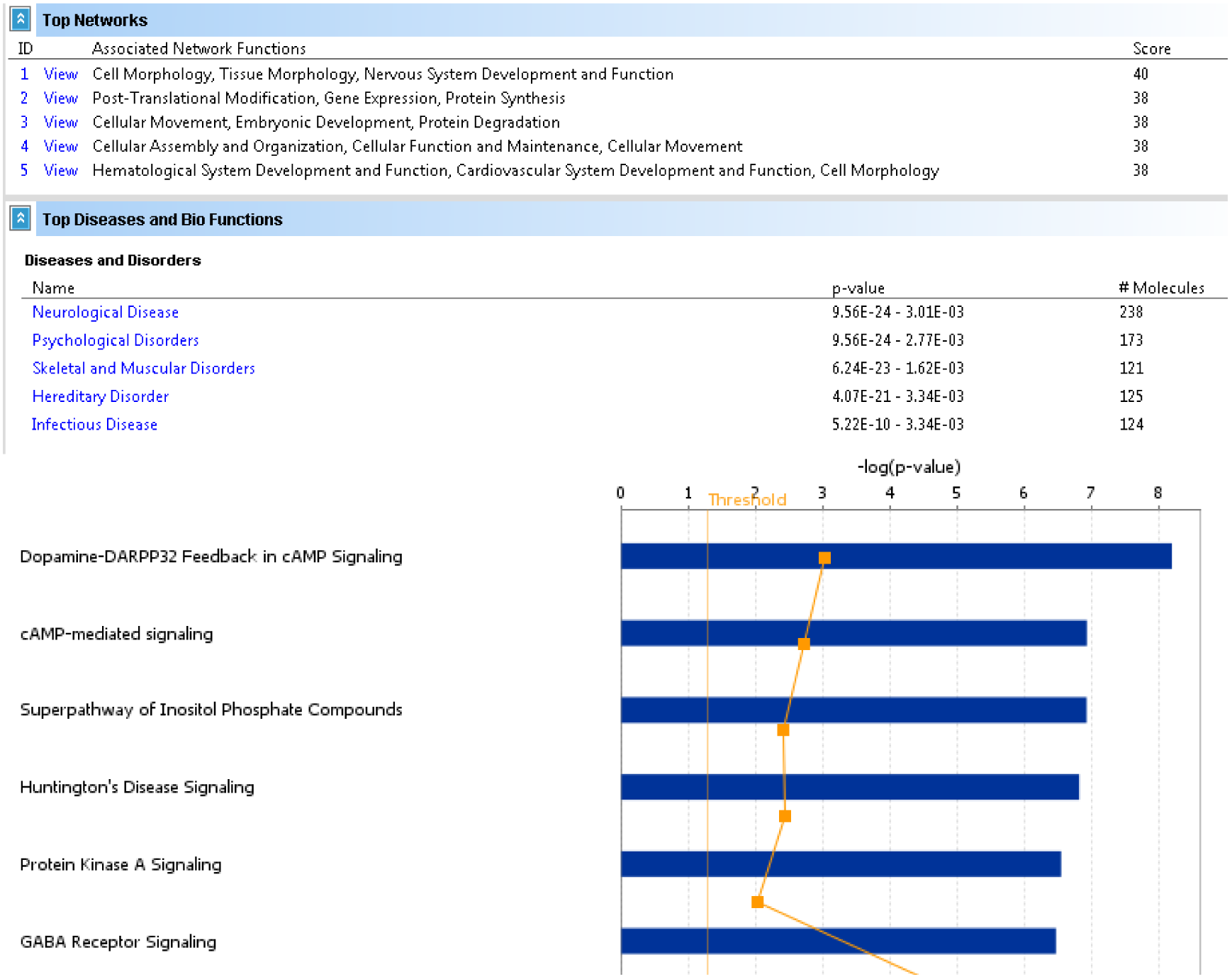

### Grade 2 group (neuron 294 probe sets)

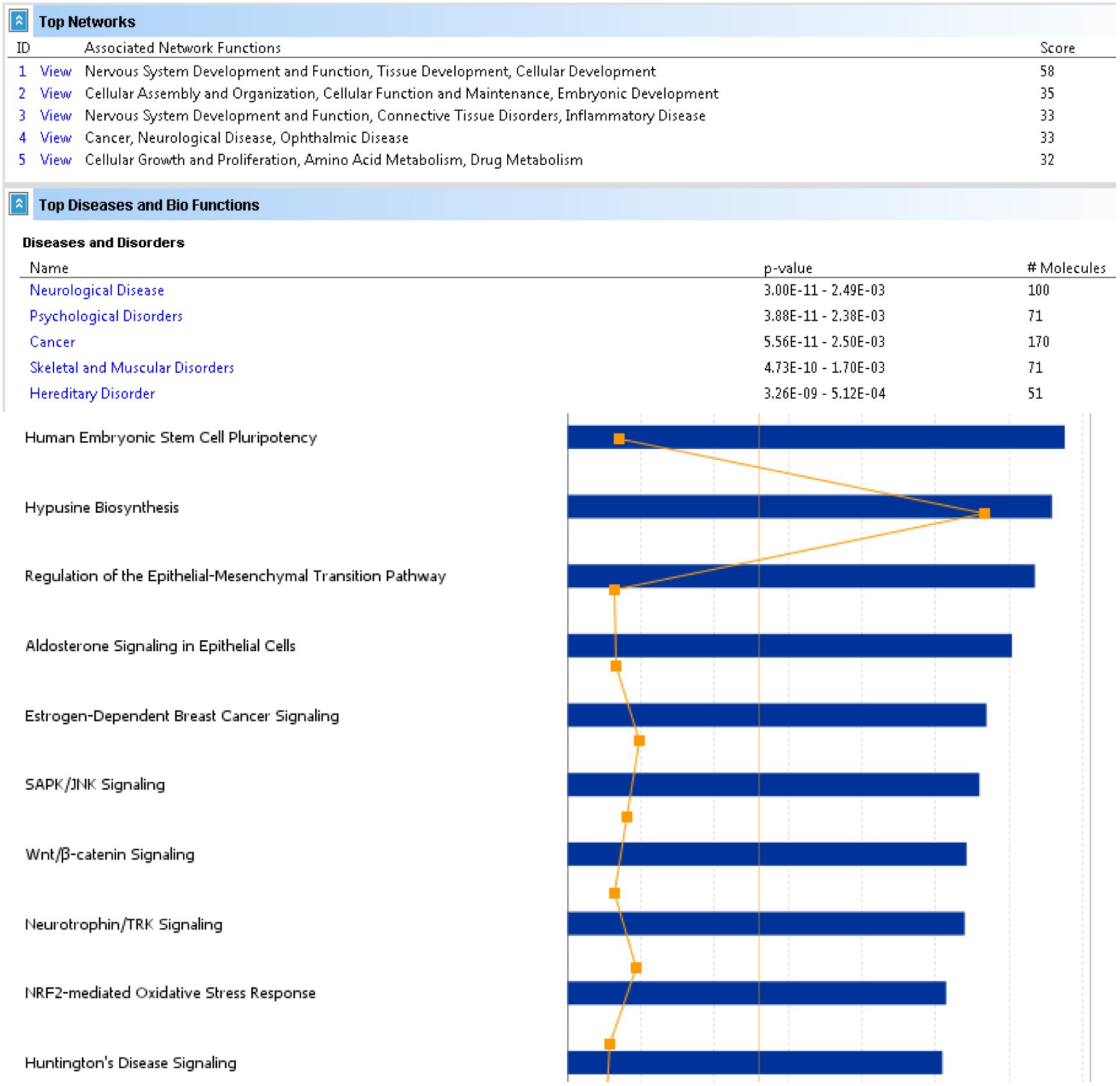

### Grade 2 group (glia 1082 probe sets)

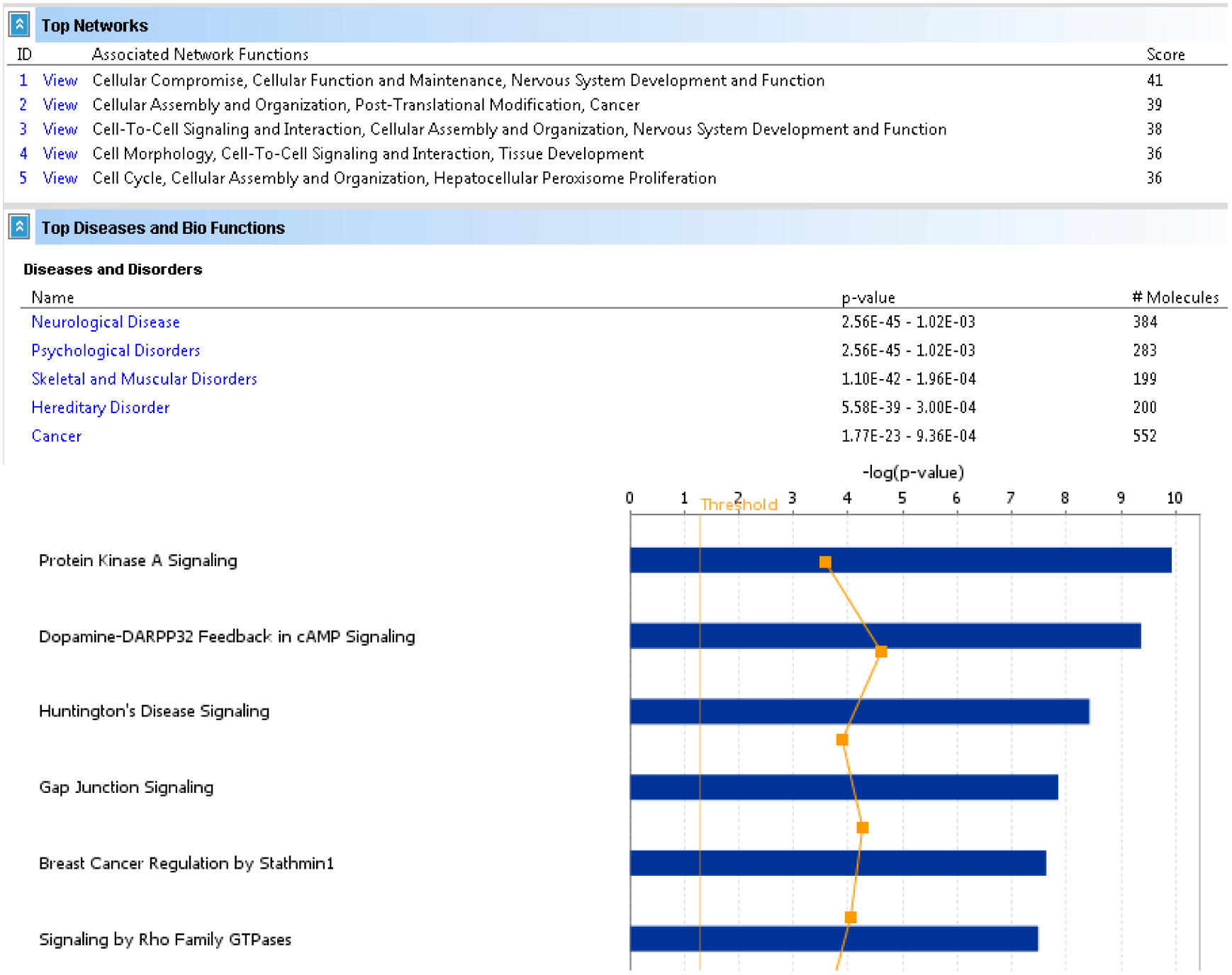

